# Implantation and Gastrulation Abnormalities Characterize Early Embryonic Lethal Mouse Lines

**DOI:** 10.1101/2020.10.08.331587

**Authors:** Yeonsoo Yoon, Joy Riley, Judith Gallant, Ping Xu, Jaime A. Rivera-Pérez

**Affiliations:** Department of Pediatrics, Division of Genes and Development, University of Massachusetts Medical School, Worcester, MA. 01655, USA

**Keywords:** Development, Mammalian, Mouse, Embryo, Gastrulation, Implantation, Knockout, Mutant, Phenotype, IMPC

## Abstract

The period of development between the zygote and embryonic day 9.5 in mice includes multiple developmental milestones essential for embryogenesis. The preeminence of this period of development has been illustrated in loss of function studies conducted by the International Mouse Phenotyping Consortium (IMPC) which have shown that close to one third of all mouse genes are essential for survival to weaning age and a significant number of mutations cause embryo lethality before E9.5. Here we report a systematic analysis of 21 pre-E9.5 lethal lines generated by the IMPC. Analysis of pre- and post-implantation embryos revealed that the majority of the lines exhibit mutant phenotypes that fall within a window of development between implantation and gastrulation with few pre-implantation and no post-gastrulation phenotypes. Our study provides multiple genetic inroads into the molecular mechanisms that control early mammalian development and the etiology of human disease, in particular, the genetic bases of infertility and pregnancy loss. We propose a strategy for an efficient assessment of early embryonic lethal mutations that can be used to assign phenotypes to developmental milestones and outline the time of lethality.

## Introduction

The period of development between the zygote and the pharyngula stage at embryonic day 9.5 (E9.5) encompasses multiple developmental milestones that are essential for survival of the embryo to term. These include cleavage, blastocyst formation, implantation, axial specification, gastrulation and establishment of the anlagen of multiple organ systems (Kojima et al., 2014; Rivera-Perez, 2013). This is also the time when crucial events like zygote genome activation (Jukam et al., 2017; Sha et al., 2019; Svoboda et al., 2015; Tadros and Lipshitz, 2009; Vastenhouw et al., 2019) and X-chromosome inactivation (Deng et al., 2014; Galupa and Heard, 2018) occur. In addition, this is the period of development when totipotent and pluripotent cells are allocated to specific pathways of differentiation. For example, segregation of trophoblast, epiblast and primitive endoderm lineages takes place between the 4^th^ and 6^th^ cell divisions (Kojima et al., 2014; Morris et al., 2010; Rossant, 2016, 2018). The prominence of the early stages of embryogenesis in human gestation is underscored by clinical studies indicating that 30% of human conceptions are lost before the embryo implants in the uterine wall (Macklon et al., 2002; Moore and Persaud, 2003). Moreover, measurements of chorionic gonadotropin, a marker of implantation, show that an additional 30% of pregnancies are lost shortly after implantation (Macklon et al., 2002). Clinical studies have also shown that the risk of early pregnancy loss increases with delayed implantation (Wilcox et al., 1999).

In mice, the significance of the early stages of development for mammalian embryogenesis has been unveiled in loss of function studies conducted by the International Mouse Phenotyping Consortium (IMPC). The IMPC is a global coalition of research institutions that aims to generate and characterize 20,000 individual knockout mouse lines (www.mousephenotype.org). A pivotal outcome of the IMPC efforts is that close to one third of all mouse genes are essential for survival to weaning age (Dickinson et al., 2016). Moreover, analysis of 242 lethal lines showed that the majority of these mutations (60.7%) are lethal before E12.5 and a large percentage of these pre-E12.5 lethal mutations (72.8%), have ceased development before E9.5 (Dickinson et al., 2016). Therefore, understanding the causes of embryo lethality at pre-E9.5 stages can provide a plethora of information on the genes that play key roles in early embryogenesis and about the developmental constraints that are responsible for the etiology of pregnancy loss and developmental abnormalities.

In this study, we aimed to elucidate the morphological anomalies of pre-E9.5 lethal lines that prevent zygote advancement through the onset of organogenesis, specifically, to assign phenotypes to developmental milestones and outline the time of lethality. Because of the possibility of maternal rescue (Bultman et al., 2006; Dodge et al., 2004; Jedrusik et al., 2015; Kim et al., 2016; Li et al., 2010), we hypothesized that the majority of the lethal phenotypes would become evident after the blastocyst stage (E3.5). As expected, only a few of the lines analyzed showed phenotypes that were evident at the blastocyst or earlier stages (4/21) while the majority of the mutant embryos (17/21) showed phenotypes between implantation and gastrulation. Intriguingly, no mutants were found to undergo normal gastrulation and reach early organogenesis stages.

These results indicate that the majority of embryonic lethal phenotypes taking place before E9.5 fall between implantation and gastrulation and suggests that maternal components may rescue pre-implantation phenotypes. The absence of post-gastrulation mutant phenotypes may reflect previous detection of this class of mutations in phenotypic screens conducted at E9.5 by the IMPC and suggests that mutant embryos that can undergo gastrulation, will survive to E9.5 or later stages.

We propose that an effective way to narrow down the phenotype of early lethal mouse lines should include an assessment of embryos at E9.5 to detect post-gastrulation abnormalities and a subsequent analysis of embryos at E3.5 to segregate lines with pre-implantation and post-implantation phenotypes. These latter phenotypes will fall between E3.5 and E7.5, a period of development encompassing implantation, egg cylinder formation, axial specification and gastrulation. Further analysis at these stages should delineate the phenotype in more detail. This strategy can outline the phenotype of early developmental mutants and guide future avenues of mammalian research.

## Results

### Pre-E9.5 mutant embryos show abnormalities at gastrulation or earlier stages

A total of 21 knockout lines containing pre-E9.5 lethal alleles were obtained from IMPC centers (**Supp. Table 1**). Each line was chosen randomly to avoid bias in specific molecular, cellular or developmental processes. We began with an assessment of E7.5 embryos derived from heterozygous crosses (**Fig. 1A**). We chose this period of development because at this time, the embryos have reached gastrulation stages and are relatively easy to assess morphologically under a dissecting microscope. For each litter, we documented the number of decidua and resorption sites. Lines that did not produce mutant embryos at E7.5 were assessed at blastocyst stages (E3.5) and some of them were subjected to embryo explant assays to determine the phenotype in more detail. We also analyzed mutant embryos at E2.5 for the lines that did not produce mutants at E3.5 (**Fig. 1A**).

**Figure 1.**
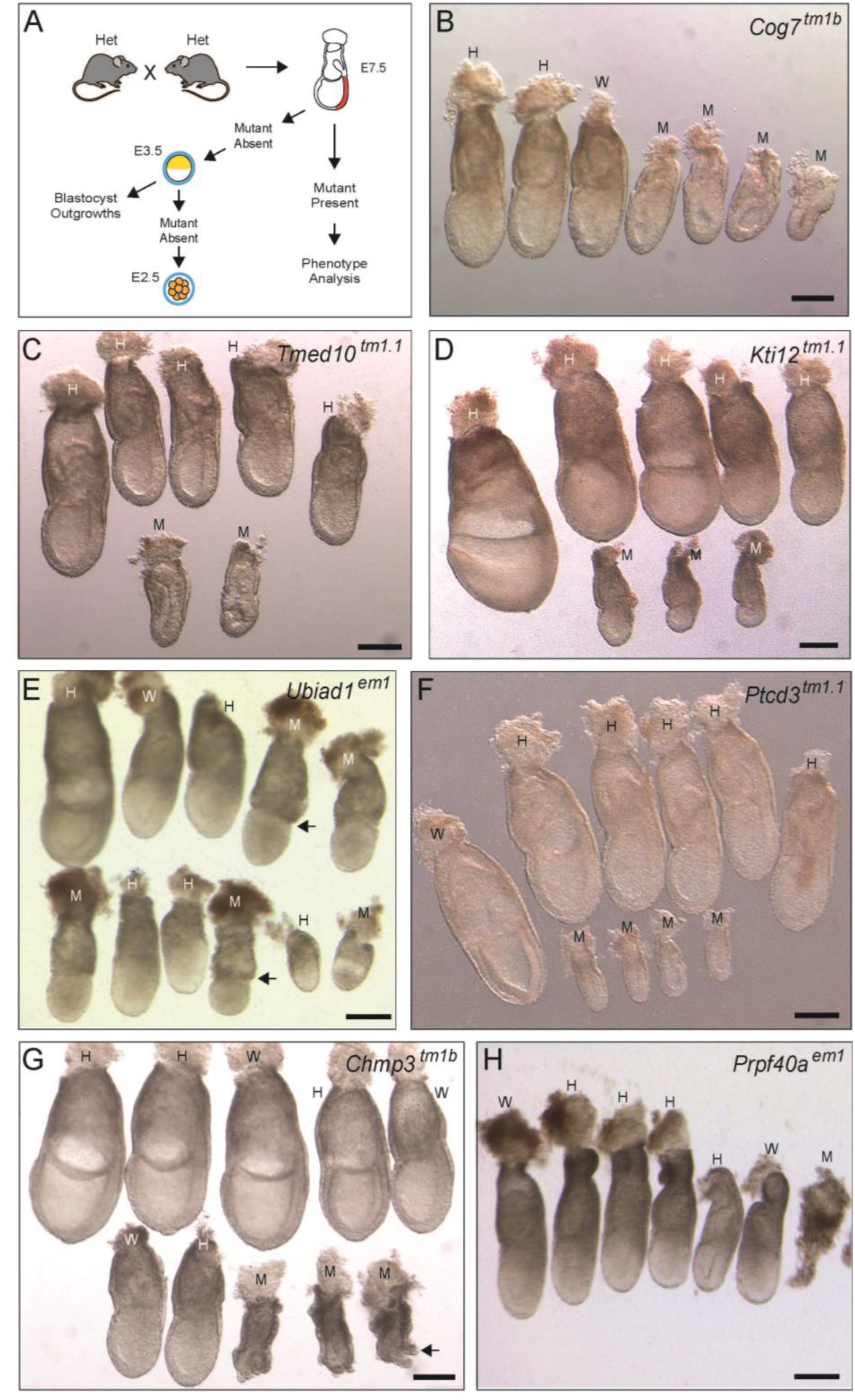
Analysis of Pre-E9.5 Lethal lines at E7.5. **A.** Schematic representation of the strategy utilized to assess the phenotype of mutant embryos. Embryos derived from heterozygous crosses were initially assessed at E7.5. For lines in which no homozygous mutants were recovered at E7.5, we analyzed embryos at E3.5 and those that did not produce mutant embryos at E3.5, were assessed at E2.5. Explant culture assays of E3.5 embryos were used to assess their ability to hatch and to form embryo outgrowths. **B-H.** Images from lines that produced mutant embryos that survived to E7.5. Mutant embryos (M), heterozygous (H) and wild-type embryos (W) are indicated. Arrows in E indicate the location of a constriction at the boundary between the epiblast and the extra-embryonic ectoderm observed in *Ubiad1*^*em1*^ mutant embryos. Arrow in G, indicates visceral endoderm bulges present in *Chmp3*^*tm1b*^ mutant embryos. Scale bars, 200 μm.

In our initial assessment at E7.5, we recovered mutant embryos from 7 lines: *Chmp3, Cog7, Kti12, Prpf40a, Ptcd3, Tmed10,* and *Ubiad1*, whereas the rest of the lines produced empty deciduae, deciduae containing remnants of resorbing embryos or did not leave any traces of mutant embryos (**Table 1**).

**Table 1.** Analysis of Pre-E9.5 Lethal lines at E7.5

| Gene / Allele | No. of Plugged Females | Not Pregnant | No. of Decidua | No. of Embryos | No. of Resorptions | Embryo Genotyping | | | $\chi^2$<br>P value |
| --- | --- | --- | --- | --- | --- | --- | --- | --- | --- |
|  |  |  |  |  |  | Mut | Het | Wt |  |
| *Alg11<sup>tm1.1</sup>* | 6 | 1 | 37 | 30<sup>a</sup> | 8 | 0 | 21 | 9 | 0.00293 |
| *Aurkaip1<sup>em1</sup>* | 5 | 2 | 30 | 31<sup>a</sup> | 0 | 0 | 19 | 12 | 0.00436 |
| *Chmp3<sup>tm1b</sup>* | 7 | 2 | 47 | 38 | 9 | 8 | 20 | 10 | 0.85394 |
| *Cog7<sup>tm1b</sup>* | 6 | 1 | 38 | 35 | 3 | 9 | 19 | 7 | 0.78438 |
| *ExoC2<sup>em1</sup>* | 5 | 0 | 50 | 40 | 10 | 0 | 24 | 16 | 0.00075 |
| *Kti12<sup>tm1.1</sup>* | 10 | 6 | 32 | 28 | 4 | 6 | 18 | 4 | 0.27645 |
| *Nup153<sup>em1</sup>* | 3 | 0 | 35 | 31 | 4 | 0 | 15 | 16 | 0.00026 |
| *Orc1<sup>tm1b</sup>* | 5 | 0 | 47 | 31 | 16 | 0 | 23 | 8 | 0.00337 |
| *Pibf1<sup>tm1.1</sup>* | 7 | 0 | 48 | 43 | 5 | 0 | 35 | 8 | 0.00005 |
| *Prpf40a<sup>em1</sup>* | 10 | 6 | 33 | 29 | 4 | 3 | 22 | 4 | 0.01996 |
| *Ptcd3<sup>tm1.1</sup>* | 6 | 1 | 44 | 40 | 4 | 8 | 22 | 10 | 0.74082 |
| *Psmc6<sup>tm1b</sup>* | 8 | 3 | 31 | 30<sup>a</sup> | 2 | 0 | 24 | 6 | 0.00136 |
| *Rfc1<sup>tm1b</sup>* | 7 | 3 | 40 | 33 | 7 | 0 | 22 | 11 | 0.00409 |
| *Rgp1<sup>tm1.1</sup>* | 7 | 1 | 56 | 35<sup>b</sup> | 21 | 0 | 25 | 9 | 0.00214 |
| *Snrnp27<sup>tm1b</sup>* | 4 | 0 | 35 | 30 | 5 | 0 | 21 | 9 | 0.0061 |
| *Srbd<sup>em1</sup>* | 8 | 3 | 45 | 30 | 15 | 0 | 22 | 8 | 0.00452 |
| *Srp9<sup>tm1.1</sup>* | 7 | 2 | 40 | 28 | 12 | 0 | 21 | 7 | 0.00525 |
| *Stx18<sup>tm2b</sup>* | 10 | 4 | 54 | 42<sup>b</sup> | 12 | 0 | 32 | 9 | 0.00026 |
| *Tmed10<sup>tm1.1</sup>* | 8 | 3 | 34 | 31 | 3 | 3 | 22 | 6 | 0.04899 |
| *Tomm20<sup>em1</sup>* | 8 | 2 | 55 | 36<sup>a</sup> | 20 | 0 | 27 | 9 | 0.00117 |
| *Ubiad1<sup>em1</sup>* | 6 | 3 | 30 | 29 | 1 | 8 | 15 | 6 | 0.85627 |

<sup>a</sup>Two embryos were found in the same decidua.
<sup>b</sup>One embryo lost, not genotyped.

These results indicate that all the lines analyzed have phenotypes that are evident at gastrulation or at earlier stages and that mutant embryos from two thirds of the lines (14/21) do not reach gastrulation. Interestingly, we did not observe mutant embryos with normal gastrulation that would indicate a post-gastrulation phenotype.

### Mutant embryos recovered at E7.5 have peri-gastrulation defects

The phenotypes of mutant embryos that survived to E7.5 varied from line to line but were fully penetrant. Mutant embryos can be classified into 5 categories: abnormal gastrulation, absence of primitive streak, developmental delay, extra-embryonic tissue abnormalities and gross morphological malformations.

Embryos exhibiting gastrulation defects were the most developmentally advanced. They initiated gastrulation, as indicated by the presence of the primitive streak, but had gastrulation abnormalities (**Fig. 1B-D**). *Cog7*^*tm1b*^ mutant embryos were smaller than littermates and appeared to have advanced to mid-streak stage (**Fig. 1B**). The posterior amniotic fold was present in some mutants but did not fuse with the anterior amniotic fold to form the amnion. Similarly, *Tmed10*^*tm1.1*^ mutants were smaller than littermates and had reached a stage close to mid-streak, but contrary to *Cog7*^*tm1*^ mutant embryos, did not show evidence of the posterior amniotic fold, suggesting an earlier gastrulation phenotype (**Fig. 1C**). *Kti12*^*tm1.1*^ mutants showed only an incipient primitive streak and were significantly smaller than littermates, reaching only about one third of the proximodistal length of the egg cylinder shown by controls (**Fig. 1D**).

Mutants for *Ubiad1*^*em1*^ did not show evidence of a primitive streak (**Fig. 1E**). They had a cylindrical epiblast morphology as opposed to an oblong cylinder shape, indicating elongation along the anteroposterior axis. A clear constriction was evident at the boundary between the epiblast and extra-embryonic ectoderm (**Fig. 1E**). *Ubiad1*^*em1*^ mutants were in general smaller than control littermates but some of them were of similar or larger size.

*Ptcd3*^*tm1.1*^ mutants were significantly smaller than control littermates, spanning approximately one quarter of the length of the proximo-distal axis of the egg cylinder of control embryos (**Fig. 1F**). There was no evidence of primitive streak formation and their small size and relative normal morphology was reminiscent of ~E6.0 embryos, suggesting developmental delay.

*Chmp3*^*tm1b*^ mutants showed extra-embryonic abnormalities. They were smaller than littermate controls and had a very characteristic phenotype consisting of bulging visceral endoderm cells (**Fig. 1G**). The most severe phenotype at E7.5 was observed in *Prpf40a*^*em1*^ mutants. These embryos consisted of small amorphous pieces of tissue surrounded by maternal components that were close to complete resorption (**Fig. 1H**).

In summary, mutant embryos that survived to E7.5 have a variety of peri-gastrulation abnormalities which suggest that attainment of gastrulation is a major barrier for subsequent embryo development.

### The majority of the mutant embryos survive to E3.5

Our initial analysis revealed that two thirds of the mutant lines (14/21) did not generate mutant embryos that survived to E7.5 (**Table 1**). To further characterize the phenotype of these lines, we analyzed embryos at E3.5. At this time of development, embryos range from compacted morulae to expanded blastocysts. This analysis allowed us to assess embryo survival during pre-implantation stages and to narrow the window of lethality.

With the exception of *Nup153*^*em1*^ and *Pibf1*^*tm1.1*^, twelve lines generated mutant embryos that survived to E3.5 (**Table 2**). A few *Nup153*^*em1*^ (1/27) and *Pibf1*^*tm1.1*^ (2/27) mutant embryos were obtained at E2.5 (**Table 2**), suggesting a requirement for these genes at cleavage or earlier stages.

**Table 2.**
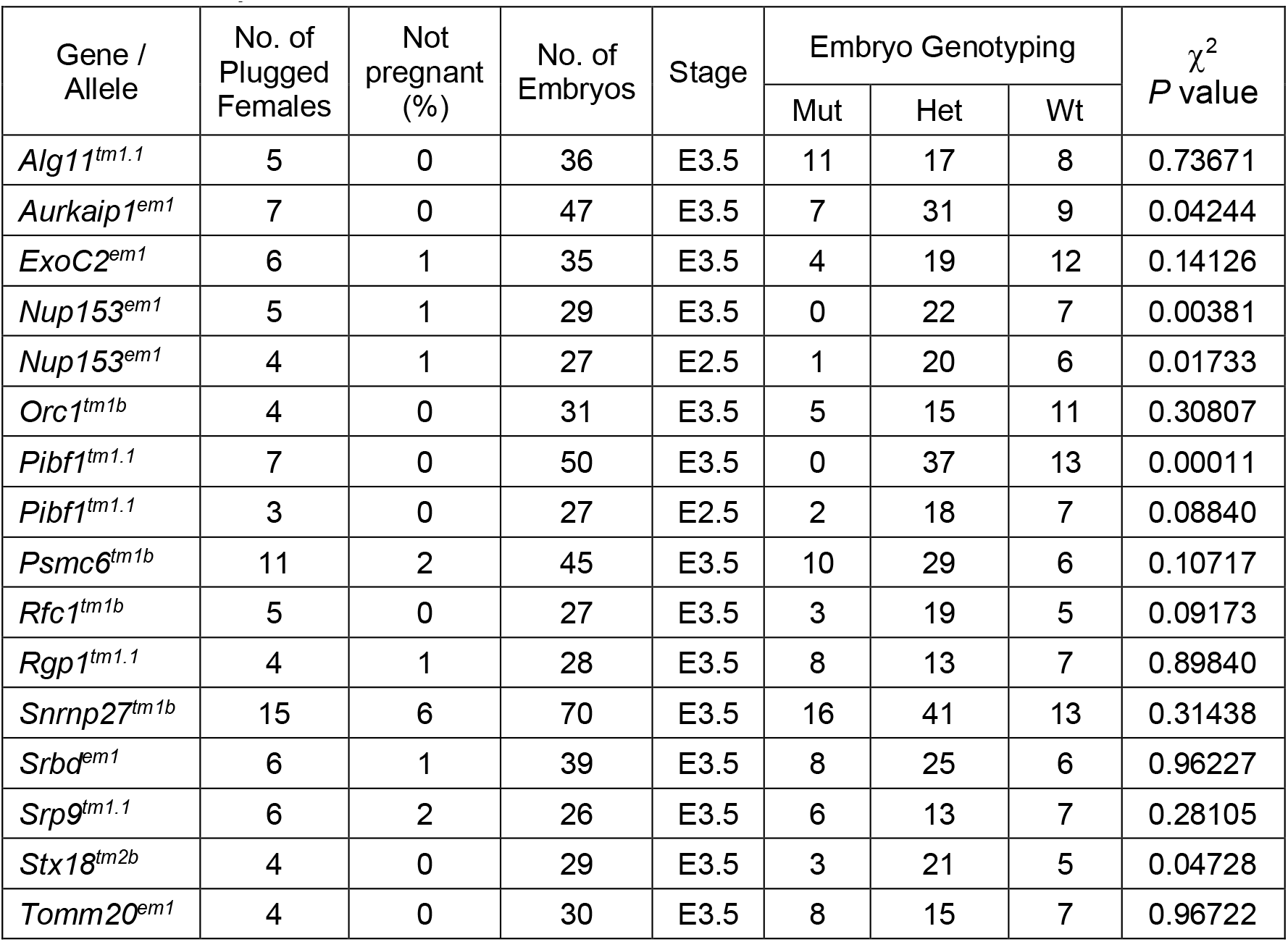
Analysis of Pre-E9.5 Lethal lines at E3.5 and E2.5

Analysis of embryo morphology revealed abnormal development of *Srbd1*^*em1*^ and *Snrnp27*^*tm1b*^ mutant embryos at E3.5. *Srbd1*^*em1*^ mutants gave rise to globular structures that resembled compacted morulae and contained cellular debris in the periphery (**Fig. 2A**).

**Figure 2.**
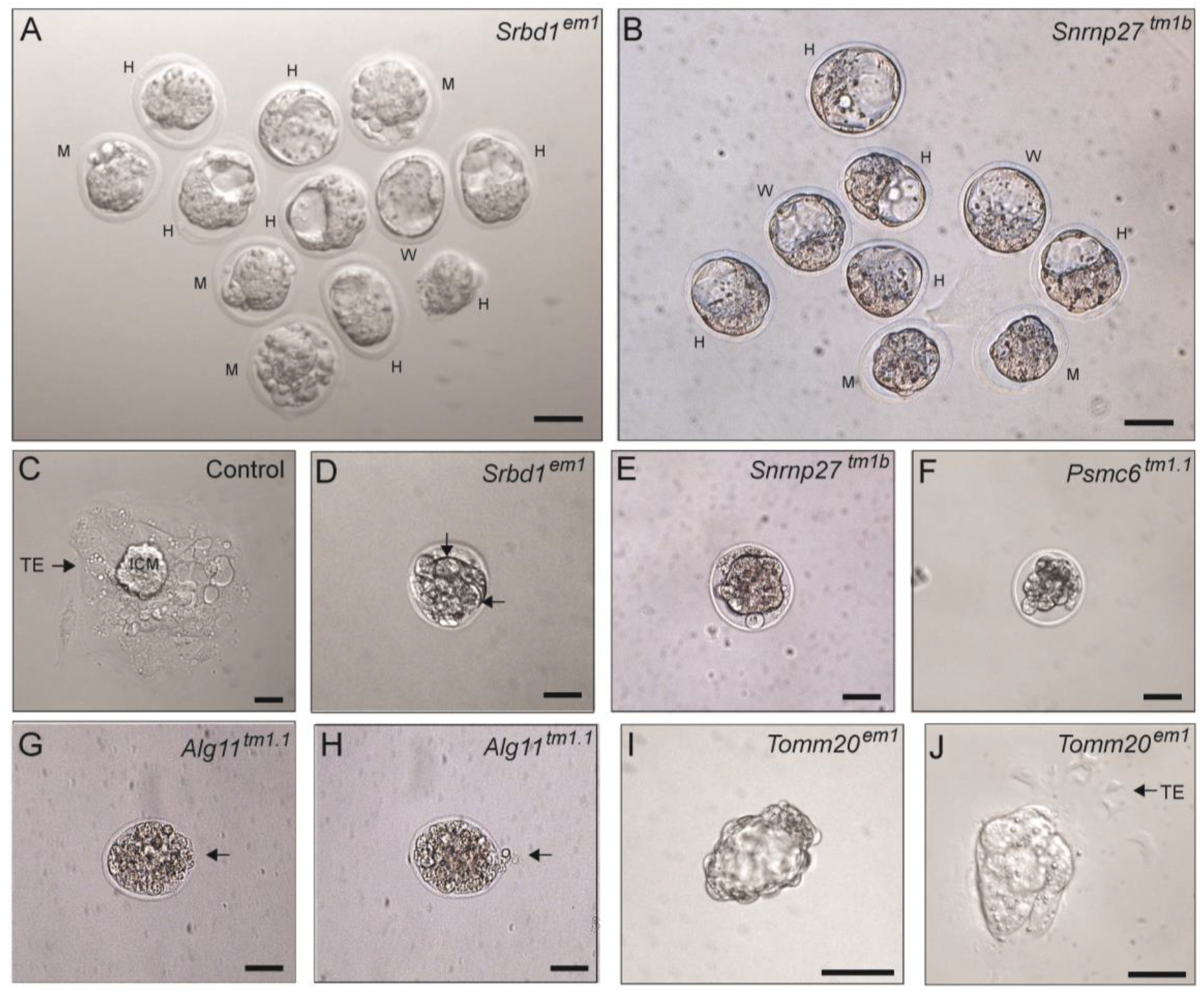
Characterization of pre-gastrulation lethal lines at E3.5. **A and B.** Litters recovered from *Srbd*^*em1*^ (**A**) and *Snrnp27*^*tm1b*^ (**B**) heterozygous crosses. Mutant embryos (M), heterozygous (H) and wild-type embryos (W) are indicated. *Srbd*^*em1*^ mutant embryos have cellular debris in the periphery. *Snrnp27*^*tm1b*^ mutant embryos were less developed than control littermates. **C.** Image of control blastocyst outgrowth cultured for four days. The outgrowth consists of an inner cell mass (ICM) derivative located in the center of the outgrowth and trophectoderm (TE) derivatives in the periphery. **D-H.** Explant cultures showing *Srbd1*^*em1*^ (**D**), *Snrnp27*^*tm1b*^ (**E**), *Psmc6*^*tm1.1*^ (**F**) and *Alg11*^*tm1.1*^ (**G, H**) mutant embryos that failed to hatch. Large vacuoles in *Srbd1*^*em1*^ embryos are indicated by black arrows. In *Alg11*^*tm1.1*^ mutants, zona erosion and partial extrusion of embryo is evident (arrows). **I and J**. *Tomm20*^*em1*^ mutant explants hatched, but failed to develop further in explant culture, forming hollow vesicles that were similar to blastocysts (**I**) or that produced underdeveloped outgrowths with poor trophectoderm development (**J**). Scale bars, 50 μm.

*Snrnp27*^*tm1b*^ mutants were at a compacted morula or early blastocyst stages and none had reached an expanded blastocyst stage (**Fig. 2B**), suggesting delayed development. The rest of the lines (10/12) did not show abnormal morphology at E3.5.

These results indicate that most of the embryos that do not survive to E7.5 can undergo cleavage and survive to E3.5, hence, the majority of the pre-E9.5 mutant lines have lethal phenotypes between E3.5 and E7.5, a period of development that encompass implantation and gastrulation.

### Explant culture experiments reveal hatching and abnormal growth phenotypes

To characterize further the phenotype of mutant embryos that survived to E3.5, we conducted explant culture assays. Explanted embryos grown in ES cell media hatch from the zona pellucida and generate outgrowths composed of trophectodermal cells attached to the bottom of the plate and a centrally located group of cells derived from the inner cell mass (**Fig. 2C**). Using this assay, we were able to evaluate mutant embryos from five lines: *Srbd1*^*em1*^, *Snrnp27*^*tm1b*^ and *Psmc6*^*tm1.1*^, *Alg11*^*tm1.1*^ and *Tomm20*^*em1*^.

*Srbd1*^*em1*^, *Snrnp27*^*tm1b*^ and *Psmc6*^*tm1.1*^ mutant embryos failed to hatch (**Fig. 2D-F**). After four days in culture, these embryos were still enclosed by the zona pellucida and had a globular morphology, suggestive of arrested development. *Srbd1*^*em1*^ explants had large vesicles that got more pronounced as the embryo degenerated further (**Fig. 2D**). *Snrnp27*^*tm1b*^ mutants appeared darkened with cellular debris in the periphery (**Fig. 2E**) and *Psmc6*^*tm1.1*^ mutants seem to be fragmenting (**Fig. 2F**).

Hatching defects were also observed in explant cultures of *Alg11*^*tm1.1*^ mutant embryos, although there were signs of partial hatching that included zona pellucida erosion or partial extrusion of the embryo (**Fig. 2G, H**). Only one *Alg11*^*tm1.1*^ embryo managed to hatch but cells formed small clumps of degenerating cells (not shown).

*Tomm20*^*em1*^ mutant embryos were able to hatch, and in some cases, attached to the bottom of the plate but generated underdeveloped outgrowths with poor trophectodermal component (**Fig. 2I, J**).

These experiments suggest that mutations in these genes disrupt trophectoderm function, which in turn, could lead to implantation failure or embryo lethality soon after implantation.

## Discussion

Our understanding of the genetic mechanisms that control development in mice has traditionally relied on a heterogeneous multipronged analysis of naturally occurring mutations as well as assessment of mutant mice generated using gene targeting strategies (Eppig et al., 2015). However, a systematic approach to characterize mouse phenotypes is highly valuable since the analysis of mouse mutations underpins our understanding of mammalian gene function and of genes linked to human disease (de Angelis et al., 2015; Dickinson et al., 2016). In this study, we analyzed pre-E9.5 lethal lines that emphasize the inactivation of a critical exon or complete deletion of a gene to ensure a full knockout (www.mousephenotype.org/understand/the-data/allele-design/). This approach together with a rigorous maintenance of the lines in a C57Bl/6NJ genetic background, allowed an earnest comparison of the phenotypes obtained. Our goal was to assign phenotypes to developmental milestones of early embryogenesis and outline the time of lethality.

We analyzed 21 knockout lines that were randomly chosen from IMPC repositories. Analysis of these lines at E7.5 revealed that only one third (7/21) produced mutant embryos that survived the first week of embryogenesis. The mutant embryos showed a variety of peri-gastrulation abnormalities that included delayed development, gastrulation defects and extra-embryonic tissue anomalies. This suggests that attainment of gastrulation is a major hurdle during embryogenesis. Intriguingly, there were no mutant embryos that looked morphologically normal at E7.5 or that had completed gastrulation, as indicated by the elevation of the neural folds, a sign of the onset of organogenesis. One possible explanation for this observation is that mutant embryos with post-gastrulation phenotypes were able to survive to E9.5 and that they were excluded from the pool of pre-E9.5 lethal lines by previous E9.5 phenotype screens conducted by the IMPC (Dickinson et al., 2016). Alternatively, it is possible that they occur at low frequency and they were not detected in our study.

E3.5 analysis of the lines that did not generate mutant embryos at E7.5 revealed that mutant embryos from most of the lines advanced to the blastocyst stage. We, however, were not able to recover *Nup153*^*em1*^ and *Pibf1*^*tm1.1*^ mutants at E3.5. Instead, a reduced number of null embryos were obtained at the 8-cell stage (E2.5). These results suggest a requirement for these genes in oogenesis, fertilization or cleavage stages. Two lines, *Srbd1*^*em1*^ and *Snrnp27*^*tm1b*^, generated embryos at E3.5 that were distinguished by morphology or delayed in development, respectively. Nonetheless, the fact that most of the lines reached the blastocyst stage suggests that some mutant phenotypes were rescued throughout the pre-implantation period by maternal components supplied by the oocyte. This is a phenomenon that has been reported previously in mutants for *Brg1*, *Eset*, *Cdx2* and *Setdb1* (Bultman et al., 2000; Bultman et al., 2006; Dodge et al., 2004; Jedrusik et al., 2015; Kim et al., 2016). Analysis of maternal mutations should reveal the functional requirement for these genes in pre-implantation development.

Embryo explant assays conducted in five lines, revealed that four of them had hatching defects and one was able to hatch but showed poor trophectoderm growth. Hatching defects and poor trophectodermal growth in these embryos suggest abnormal trophectoderm function since this tissue controls hatching and implantation (Perona and Wassarman, 1986; Russ et al., 2000; Schiewe et al., 1995; Strumpf et al., 2005; Wang and Dey, 2006), which in turn, can lead to implantation failure or lethality soon afterwards.

Overall, our results indicate that the majority of pre-E9.5 lethal strains generate mutant embryos that can survive to the blastocyst or later stages of development up to gastrulation. In **figure 3**, we illustrate these results relative to the time of gestation and key events of embryogenesis.

**Figure 3.**
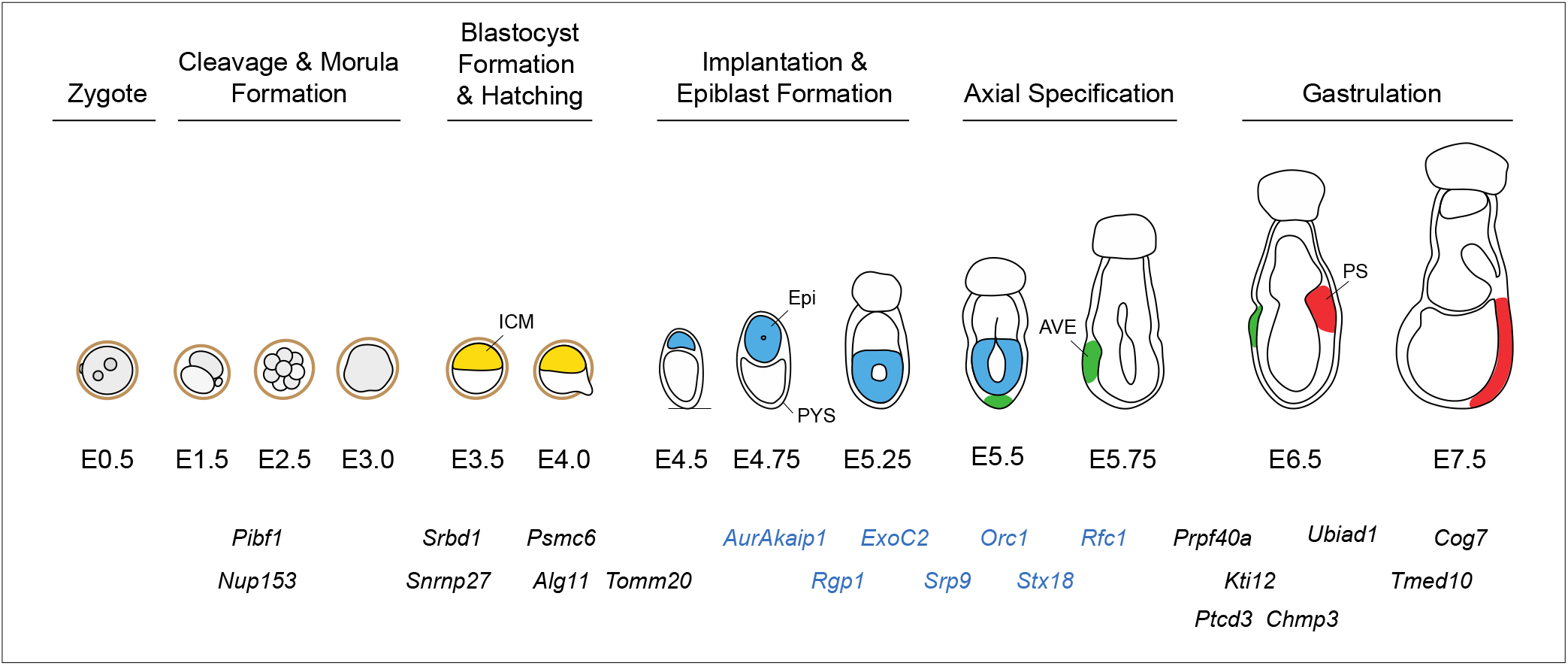
Approximate onset of lethal phenotypes relative to developmental landmarks. Mutant embryos exhibit a variety of phenotypes that span pre-implantation and gastrulation stages (E2.5-E7.5). The majority of the mutations (19/21) affected stages between the blastocyst stage and gastrulation (E3.5-E7.5). Mutants for genes marked in blue had normal blastocyst morphology but were not able to develop to E7.5, suggesting lethality between implantation and pre-gastrulation stages (E4.0-E6.5). For simplicity, the parietal yolk sac is not depicted in conceptuses from E5.25 onwards. ICM, inner cell mass (yellow); Epi, epiblast (blue); AVE, anterior visceral endoderm (green); PS, primitive streak (red) and PYS, parietal yolk sac.

Our data shows that about 50% of the lines studied (11/21) can implant and continue development over a post-implantation window of time that precedes the onset of gastrulation (~E4.0 − ~E6.5). These lines may have phenotypes that affect the formation of the egg cylinder, the process of axial specification or have defects that prevent them from advancing to gastrulation. Kojima and co-workers have suggested that attainment of a threshold number of cells has a strong correlation with formation of the proamniotic cavity in the epiblast and attainment of gastrulation (Kojima et al., 2014). This is based on blastomere reduction experiments (Power and Tam, 1993; Rands, 1986), however, analysis of tetraploid embryos which have half the cell number but increased cell volume due to hyperploidy reach gastrulation at the same time as diploid embryos (Henery et al., 1992). Therefore, it is suggested that the beginning of gastrulation may be related to tissue size not just cell number (Kojima et al., 2014). In the case of early post-implantation lethal lines, it is possible that mutations that lead to low cell numbers at pre-gastrulation stages will not reach the critical cell number to allow proper morphogenesis of the epiblast or reach the critical tissue mass to initiate gastrulation.

The majority of the lines for which mutant embryos were absent at E7.5, produced resorption sites that consisted of empty decidua or decidua that contained debris of resorbing embryos. These included lines with cleavage phenotypes like *Nup153*^*em1*^ and *Pibf1*^*tm1.1*^ for which no mutant embryos were observed at E3.5. This suggest that some pre-implantation lethal embryos can survive to post-implantation stages. However, an alternative explanation is that resorptions obtained from these lines represent heterozygous or wild-type embryos that perished after implantation. This is also the case of *Snrnp27*^*tm1b*^ embryos who were delayed in development at E3.5 and failed to hatch in explant culture experiments.

The frequency of resorption sites appeared to correlate with the severity of the phenotype. The number of resorptions amounted to 10%, 11% and 14% in *Nup153*^*em1*^, *Pibf1*^*tm1.1*^ and *Snrnp27*^*tm1b*^ mutant lines respectively, which had the earliest phenotypes. In contrast, analysis of litters from *Alg11*^*tm1.1*^ and *Tomm20*^*em1*^, which had less severe phenotypes, revealed higher proportion of resorption sites that amounted to 22% and 36%, respectively.

This suggests that the most severely affected mutants perish earlier in development and traces of the embryos or remnants of implantation, do not last to E7.5. An exception was *Srbd1*^*em1*^ mutants which led to abnormal blastocysts morphology and failed to hatch in explant culture assays, yet generated resorption sites at a frequency of 33%. These results suggest that the uterus provides a more favorable environment for embryo survival than *in vitro* culture conditions.

The data presented here, should be viewed as the starting point for future experiments that can shed light on the mechanisms of early mammalian development as well as inform us on the etiology of human disease. It has been postulated that a clock mechanism controlling pre-implantation development may be linked to titration of inhibitory cytoplasmic factors or mechanism associated with DNA replication, DNA modification and chromatin remodeling (Kojima et al., 2014). Thus, analysis of phenotypes with pre-implantation lethality and subsequent transcriptional profiling can define molecular pathways and reveal candidates that mediate cleavage and the transition through compaction and blastocyst formation. Current models suggest that a cell sorting mechanism driving the segregation of epiblast and primitive endoderm cells begins in the ICM of E3.5 blastocysts (Chazaud and Rossant, 2006; Plusa et al., 2008). Phenotypes affecting this process will be reflected in the formation of the primitive endoderm which can be revealed by markers of primitive endoderm such as *Pdgfrα* (Artus et al., 2010). Analysis of Nanog and Gata6 in the ICM will also shed light on this phenomenon (Frum et al., 2013; Rossant, 2016; Schrode et al., 2013).

Some lines showed a high proportion of plugged females that were not pregnant. These lines include *Kti12*^*tm1.1*^ and *Prpf40a*^*em1*^ in which 6 out of 10 plugged females did not produce embryos. These results suggest uterine anomalies or other defects indicative of limited fertility in heterozygous animals. This is an exciting possibility that can inform human fertility research since almost a quarter of clinical infertility cases in humans are idiopathic (Matzuk and Lamb, 2002). Further studies will be necessary to confirm these results and determine if anatomical defects, gonadal dysfunction, endocrinopathies or other defects are responsible for these fertility abnormalities (Matzuk and Lamb, 2002).

Biological systems have an inherent ability to maintain phenotypic stability shaped by evolution that allows them to thrive in an unpredictable environment (de Visser et al., 2003). Phenotypic stability, defined as the ability of a biological system to maintain its functions despite internal and external perturbations, is known as robustness (Kitano, 2004; Stelling et al., 2004). Robustness may explain the survival of 66% of the mutant lines generated by the IMPC that are homozygous viable and another 9.6% that are subviable (https://www.mousephenotype.org/data/embryo). In fact, evidence suggests that subviable lines are more likely to have a paralog gene than lethal ones (Dickinson et al., 2016). In zebrafish, genetic robustness has been proposed to occur by a process of compensation called transcriptional adaptation in which mutant mRNA degradation triggers the upregulation of genes that exhibit sequence similarity to the mutated gene (El-Brolosy et al., 2019; El-Brolosy and Stainier, 2017). Expression profiling of mutant and control embryos will allow us to explore these possibilities.

As previously mentioned, the IMPC plans to generate and functionally characterize mutations in 20,000 mouse genes and expects that a significant proportion of the mutated genes will have lethal phenotypes during early embryogenesis (www.mousephenotype.org/about-impc/). Therefore, an efficient strategy will have to be employed to functionally annotate these lethal phenotypes. Our study indicates that the majority of pre-E9.5 lethal lines exhibit phenotypes that fall within a window of development between implantation and gastrulation (E4.0 and E7.5) with a few pre-implantation and no post-gastrulation phenotypes. Hence, an efficient strategy to narrow down the phenotype of lethal mutations occurring at early stages of embryogenesis is to analyze embryos at E9.5 to detect post-gastrulation phenotypes and at E3.5 to segregate pre- and post-implantation phenotypes. Further analysis at E7.5 (or E6.75) should help distinguish phenotypes occurring at pre-gastrulation or during gastrulation stages. Additional analyses at other stages and blastocyst explant culture experiments combined with expression profiling should help pinpoint the molecular and cellular mechanisms driving the phenotypes observed.

## Acknowledgements

We thank Zdenka Matijasevic for critically reading the manuscript. This work was supported by NIH grant HD083311 to JAR-P.

## Authors Contributions

YY, dissected E7.5 embryos, conducted analysis of mutant phenotypes, performed embryo explant experiments, analyzed results and wrote the manuscript together with JAR-P; JR, Conducted analysis of mutant phenotypes, performed embryo explant experiments, gathered results, maintained mouse colony, genotyped embryos and mice; JG and PX conducted *in vitro* fertilization experiments; JAR-P, conceived and planned the experiments, dissected E7.5 embryos, analyzed data, wrote the manuscript.

## Declarations of Interests

The authors declare no competing interests.

## Materials and Methods

### Mice

All experiments were conducted under the guidance of the Institutional Animal Care and Use Committee of the University of Massachusetts Medical School. Live heterozygous mice or frozen sperm from mutant lines were obtained from repositories at the Jackson Laboratory, Baylor college of Medicine or the University of California-Davis (Supplementary Table 1). All mice were maintained in C57BL/6NJ background. To generate additional heterozygous animals, founder mice were mated to wild-type C57Bl/6JN mice. Frozen sperm was used to fertilize oocytes obtained from C57Bl/6JN females as described below. Animals were maintained on a 12 h light cycle. The mid-point of the light cycle of the day that a mating plug was observed was considered embryonic day 0.5 (E0.5) of gestation.

### *In vitro* fertilization

Females were superovulated using a HyperOva (anti-inhibin antisera) and hCG combination, as previously described (Nakagawa et al., 2016; Takeo and Nakagata, 2015). *In vitro* fertilization was conducted with a recently described method that leads to high fertilization rates from shipped cryopreserved sperm (Nakagata et al., 2014; Takeo and Nakagata, 2018). This procedure allowed us to generate large cohorts of heterozygous animals of similar age to simultaneously generate multiple embryos for phenotype analysis.

### Embryo staging

Conceptuses were staged using morphological landmarks as described by Downs and Davis (Downs and Davies, 1993) and Rivera-Perez and co-workers (Rivera-Perez et al., 2010).

Cleavage stage embryos were staged by counting the number of blastomeres. Blastocysts were staged using the ratio of polar to mural trophectoderm based on the description of Gardner (Gardner, 1997) using the following classification: very early blastocyst (3:1), early blastocyst (2:1), expanding blastocyst (1:1) and expanded blastocyst (1:2 or more).

### Explant culture assays

Embryos were collected from uteri of pregnant females at E3.5 as previously described (Behringer et al., 2014). Litters were imaged using an inverted fluorescence microscope (Leica, DMI4000). To generate outgrowths, each embryo was cultured in individual drops of DMEM containing 10% FBS for 4-7 days in a tissue culture incubator supplemented with 5% CO_2_ and 5% O_2_.

### Isolation and analysis of E7.5 embryos

E7.5 embryos were dissected as follows: heterozygous females mated with heterozygous males were euthanized by cervical dislocation. The uterine horns were removed, placed in PBS and each implantation site was dissected manually using forceps to isolate the decidua. The shape and size of each decidua was assessed to detect potential anomalies that could indicate the presence of abnormal embryos. Decidua were then transferred to DMEM media containing 20 mM HEPES. Each decidua was split longitudinally to access the embryos (Behringer et al., 2014). At this time, the parietal yolk sac was assessed for the presence of debris or blood infiltration that may indicate abnormal function or defective embryos. After removal of the parietal yolk sac, the shape and the length of the proximodistal axis of the egg cylinder (from the base of the ectoplacental cone to the tip of the epiblast) was evaluated visually and compared with littermates.

### Genotyping

The genotype of mice and E7.5, E3.5 and E2.5 embryos was determined by PCR using the oligos described in Supplementary Table 2. Mouse tails (2 mm tail tip piece) were placed in 200 μl of PCR lysis buffer (50 mM KCl, 10 mM Tris-HCl pH8.3, 2.5 mM MgCl_2_, 0.1 mg/ml gelatin, 0.45% IGEPAL and 0.45% Tween 20) containing 100 mg/ml Proteinase K and incubated overnight at 56°C. After lysis, the proteinase K was inactivated at 95°C for 8 min and 0.5 - 1 μl of the sample was used for the PCR reaction. E7.5 embryos were lysed in 20 μl and E3.5 and E2.5 embryos in 10 μl of PCR lysis buffer. The ectoplacental cone of E7.5 embryos was removed before PCR analysis to avoid maternal contamination.

### Statistical Analysis

The statistical significance of embryo frequencies obtained from heterozygous crosses was assessed using Chi-Square test (d.f.=2, α 0.05). We assessed a minimum of 26 or 28 embryos per line at E3.5 and E7.5 respectively.

---

**Supplementary Table 1.**
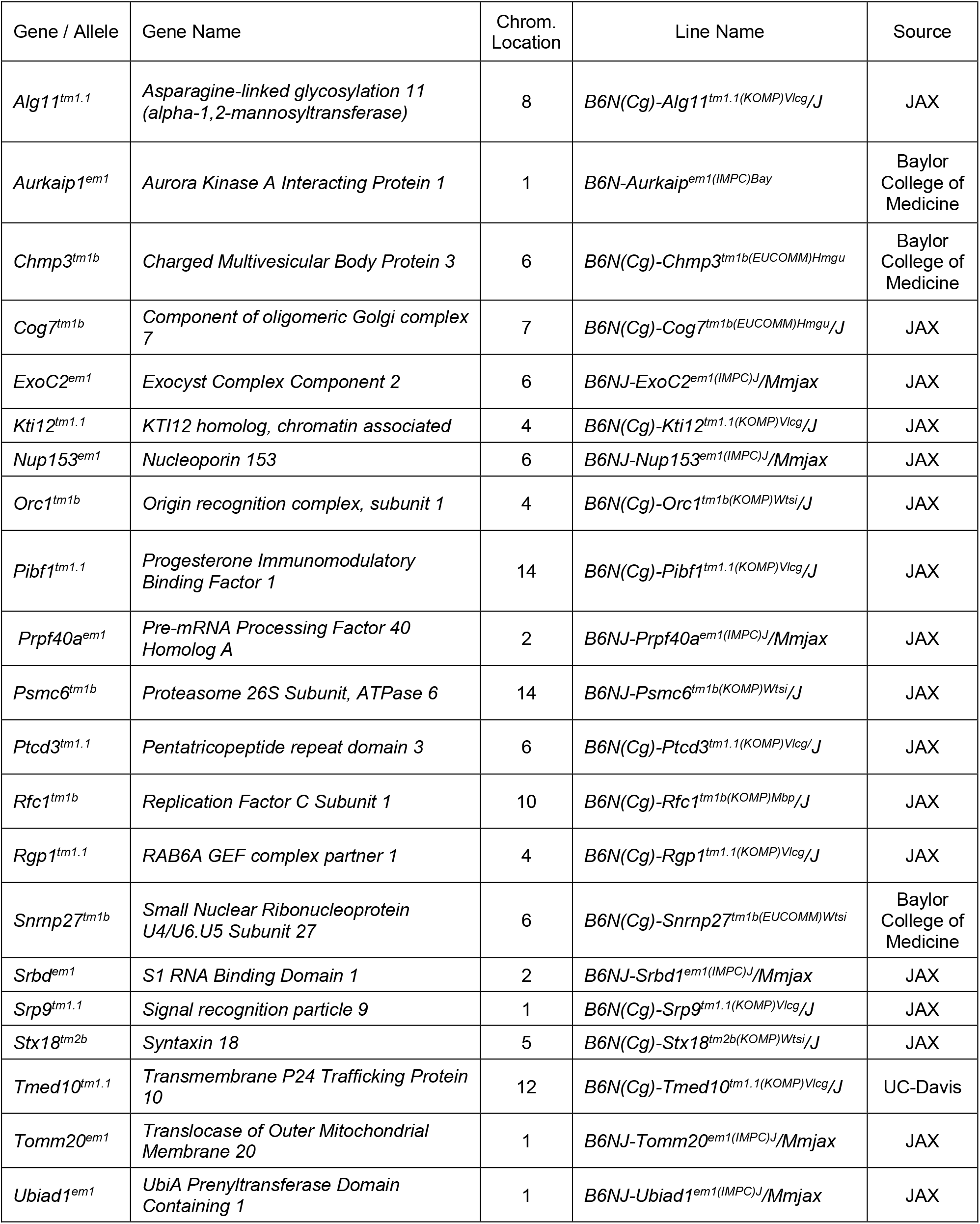
E9.5 Lethal Lines Analyzed

**Supplementary Table 2.**
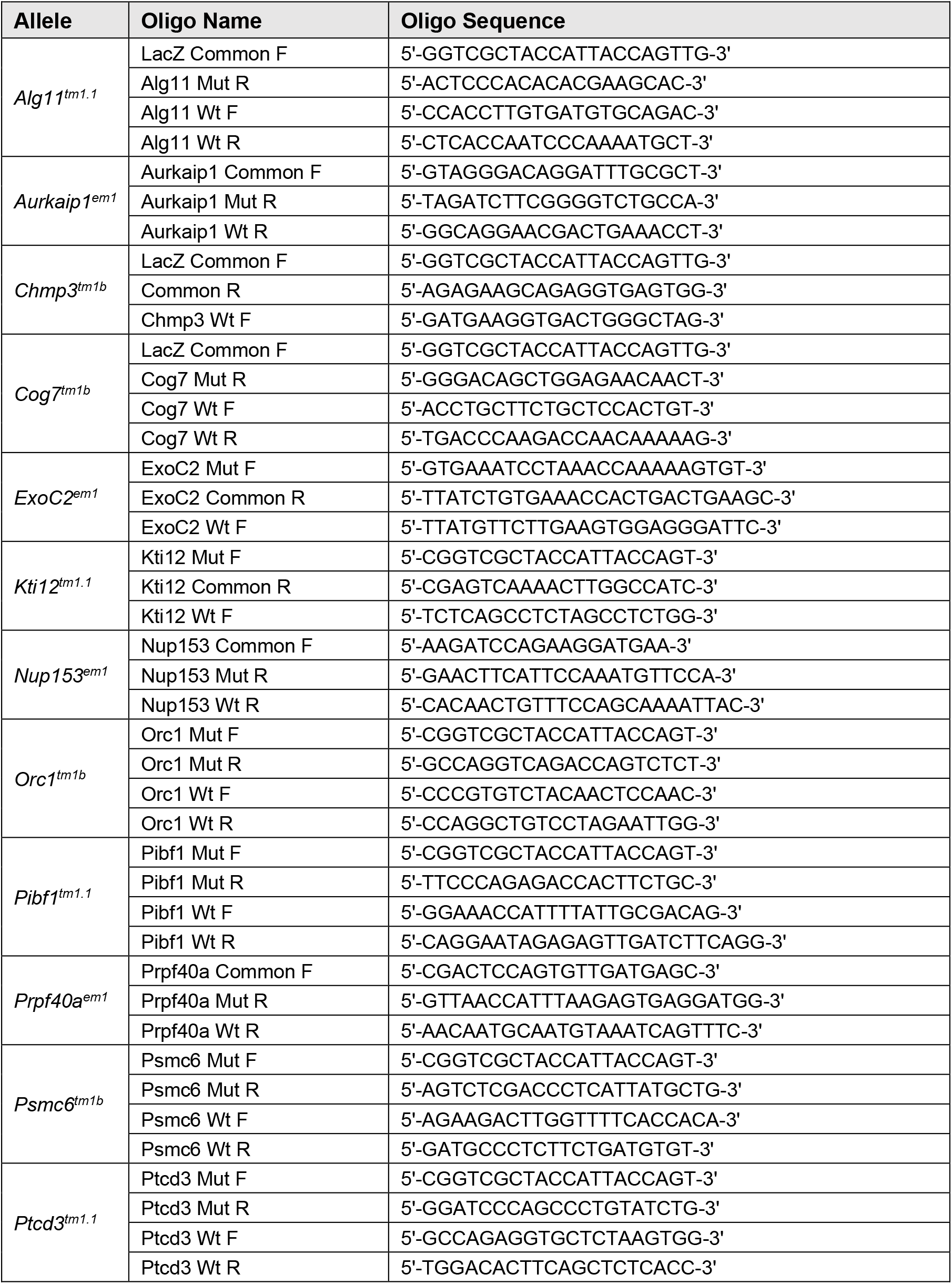
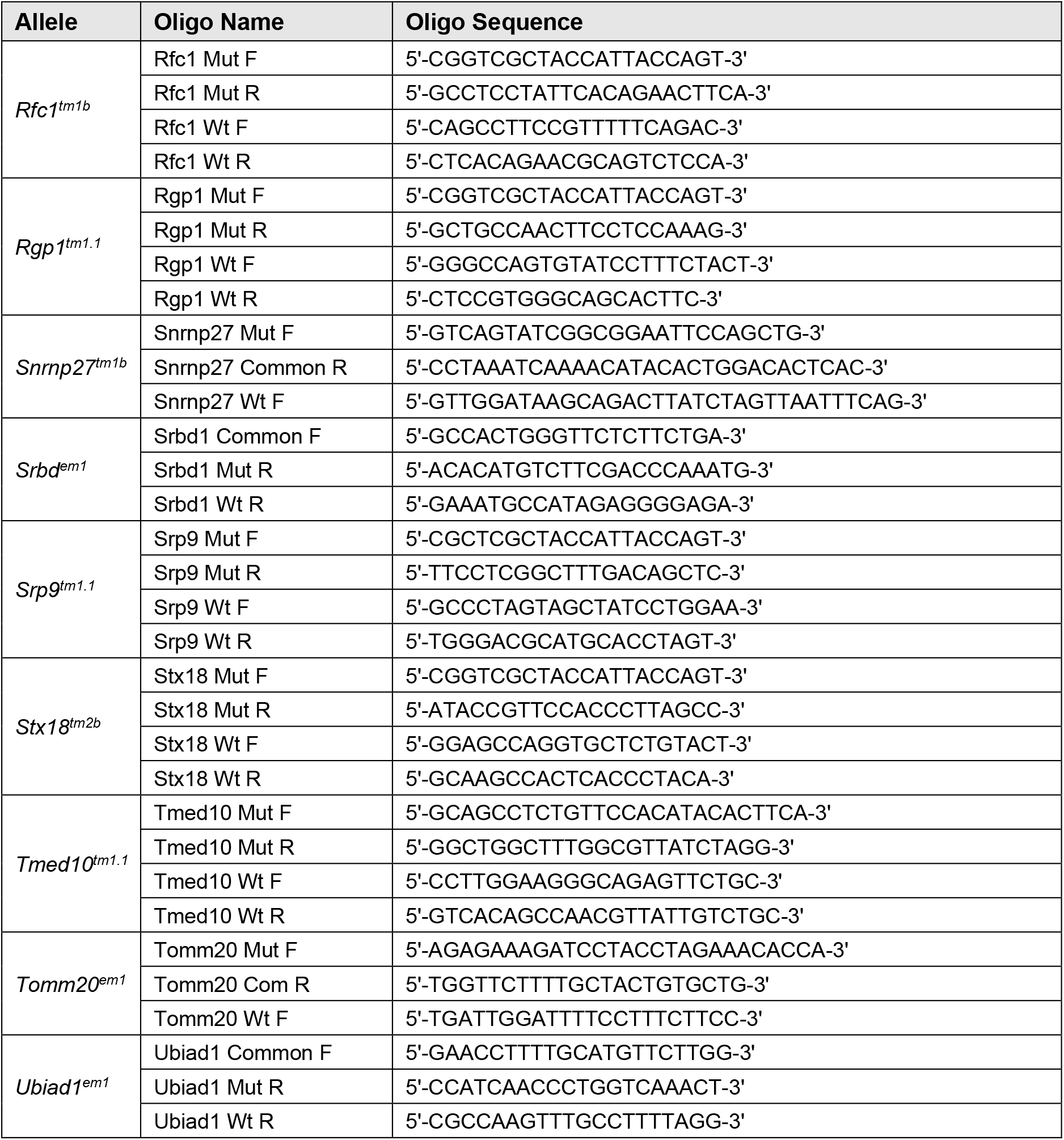
List of oligos used for genotyping

## Notes

### Competing Interest Statement

The authors have declared no competing interest.

